## Supplementary material for "Flotillin mediated membrane fluidity controls peptidoglycan synthesis and MreB movement": Figure supplements and legends

1) Supplementary Movie legends
2) Supplementary Figures with legends.
3) Supplemental Methods.

### Supplementary Movie legends.

**Supplemental Movie 1. Visualization of xylose inducible mrfpRuby-MreB patches dynamics (strain 4070) during exponential growth in LB medium at 37°C by TIRF microscopy.**

Exposure time was 2 sec and frame rate 1 image/sec over 30 seconds. MreB patches rotate perpendicularly to the longitudinal cell axis with an average speed of 92.98 nm/s (+/- 21.37). This movie refers to MreB speed presented in figure 4. Speed of the movie: 15 fps

**Supplemental Movie 2. Visualization of xylose inducible mrfpRuby-MreB patches dynamics (strain 4070) during exponential growth in SMM medium at 37°C by TIRF microscopy.**

Exposure time was 2 sec and frame rate 1 image/sec over 30 seconds. MreB patches rotate perpendicularly to the longitudinal cell axis with an average speed of 58.09 nm/s (+/- 15.08). This movie refers to MreB speed presented in figure 4. Speed of the movie: 10 fps

**Supplemental Movie 3. Visualization of xylose inducible mrfpRuby-MreB in Δ*floAT* patches dynamics (strain 4076) during exponential growth in LB medium at 37°C by TIRF microscopy.**

Exposure time was 2 sec and frame rate 1 image/sec over 30 seconds. MreB patches rotate perpendicularly to the longitudinal cell axis with an average speed of 41.59 nm/s (+/- 10.48). This movie refers to MreB speed presented in figure 4. Speed of the movie: 15 fps

**Supplemental Movie 4. Visualization of xylose inducible mrfpRuby-MreB in Δ*floAT* patches dynamics (strain 4076) during exponential growth in SMM medium at 37°C by TIRF microscopy.**

Exposure time was 2 sec and frame rate 1 image/sec over 30 seconds. MreB patches rotate perpendicularly to the longitudinal cell axis with an average speed of 65.88 nm/s (+/- 15.68). This movie refers to MreB speed presented in figure 4. Speed of the movie: 15 fps

**Supplemental Movie 5. Visualization of xylose inducible mrfpRuby-MreB patches dynamics (strain 4070) during exponential growth in LB medium supplemented with BnOH (0.1%) at 37°C by TIRF microscopy.**

Exposure time was 2 sec and frame rate 1 image/sec over 30 seconds. MreB patches rotate perpendicularly to the longitudinal cell axis with an average speed of 79.00 nm/s (+/- 22.25). This movie refers to MreB speed presented in figure 4. Speed of the movie: 15 fps

**Supplemental Movie 6. Visualization of xylose inducible mrfpRuby-MreB in Δ*floAT* patches dynamics (strain 4076) during exponential growth in LB medium supplemented with BnOH (0.1 %) at 37°C by TIRF microscopy.**

Exposure time was 2 sec and frame rate 1 image/sec over 30 seconds. MreB patches rotate perpendicularly to the longitudinal cell axis with an average speed of 69.22 nm/s (+/- 25.03). This movie refers to MreB speed presented in figure 4. Speed of the movie: 15 fps

**Supplementary Figures.**

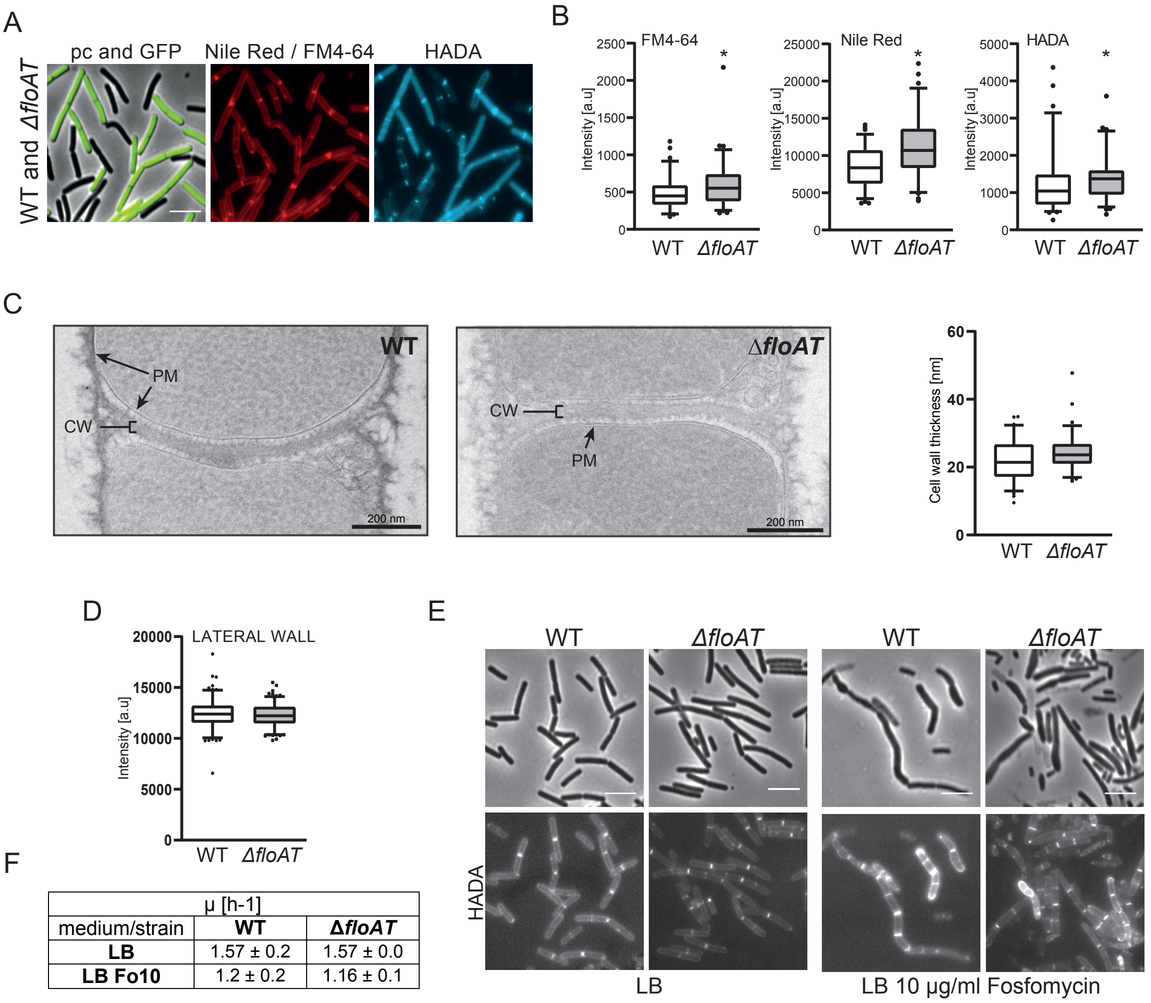


| μ [h-1] | | |
|---|---|---|
| medium/strain | **WT** | **ΔfloAT** |
| **LB** | 1.57 ± 0.2 | 1.57 ± 0.0 |
| **LB Fo10** | 1.2 ± 0.2 | 1.16 ± 0.1 |

**Figure 1 – Figure supplement 1. A**. Morphology of the exponentially growing 4092 (WT-GFP) and Δ*floAT* strains labelled simultaneously with Nile Red or FM 4-64 and HADA. **B**. Septal peak intensity of FM4-64, Nile Red and HADA labelled division sites of the cells shown in (A). Cells from each strain (n = 70) were analysed on the same agarose pad using the ObjectJ macro tool PeakFinder followed by statistical analysis with Prism resulting in box plot graphs. Significant differences are based on the two-tailed Mann-Whitney test (* p<0.05). **C.** Electron micrographs of the septal plane of the exponentially growing 168 (WT) and *ΔfloAT* strains alongside with a comparison of their cell wall thickness analysis represented as box plot graphs (n=70). **D.** Peak intensities of Nile-Red labelling of the lateral membranes of the WT and Δ*floAT* strains depicted in (A). A Mann-Whitney T-test ($p$ < 0.05, n ≥ 160), showed no significant statistical difference in intesntity between the tested strains. **E.** Morphology of exponentially growing 168 (WT) and *ΔfloAT* strains cultivated in rich (LB) medium grown with or without a sub-lethal concentration of Fosfomycin (LB + 10 µg/ml Fosfomycin) and labelled with HADA. Scale bar 5 µm. **F.** Growth rate of the cells depicted in (E) based on two biological replicates and three technical repetitions.

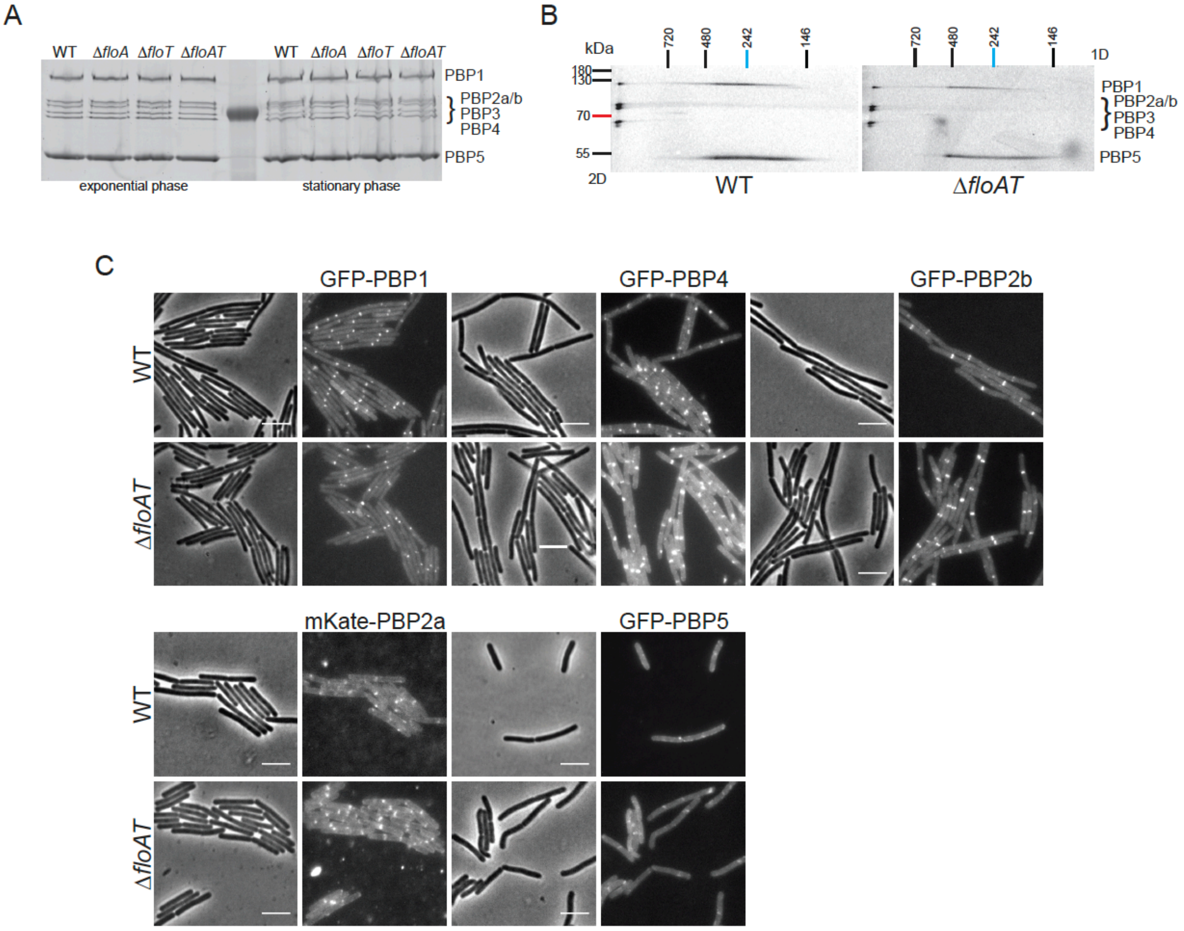


**Figure 1 – Figure supplement 2. Absence of flotillins does not affect expression, oligomerization or localization of PBPs.**

**A**. Expression pattern of PBPs in wild type and flotillin deficient strains visualized with Bocillin-FL. Membrane fractions isolated from cells (wt, Δ*floA*, Δ*floT* and Δ*floAT*) in exponential and stationary phase of growth were labelled with Bocillin-FL and run on an SDS-PAGE. The fluorescent signal was detected using Typhoon® Trio scanner. **B**. Twodimensional (2D) BN/SDS-PAGE of PBP membrane oligomers labelled with Bocillin-FL. Membrane fractions of wt and Δ*floAT* strains were isolated, labelled with Bocillin-FL and loaded onto Blue-Native PAGE. The respective lanes were excised, horizontally immobilized on top of the SDS-PAGE gel and resolved. The fluorescent signal was detected using a Typhoon® Trio scanner. PBPs 1, 2, 3 and 4 are present in a complex that is not resolved in the native gel, which is resolved at the left of the second dimension gel. PBP5 cannot be found in this complex but runs across a continuum of mass in the second dimension gel. A similar continuum is found for a second fraction of PBP1. This continuum either comes from a second complex that is disintegrating in the first dimension or from various complexes with different but close masses. Although our analysis clearly indicates that several PBPs are part of high Mw complexes, no differences in the patterns of the PBP complexes were found in membranes of the Δ*floAT* strain. **C**. Localization patterns of GFP-PBP fluorescent fusions in flotillin deletion strains. Exponentially growing wild-type (wt) and flotillin deletion (Δ*floAT*) strains expressing the indicated GFP-PBP fusions were imaged by phase contrast and fluorescence microscopy. The PBPs chosen were the two main aPBPs 1 and 4, the division associated bPBP 2b, the elongation associated bPBP 2a, and the main D,D-carboxypeptidase PBP5. No obvious differences in the localization patterns for the PBPs were detected in the absence of flotillins. Scale bar 5 µm.

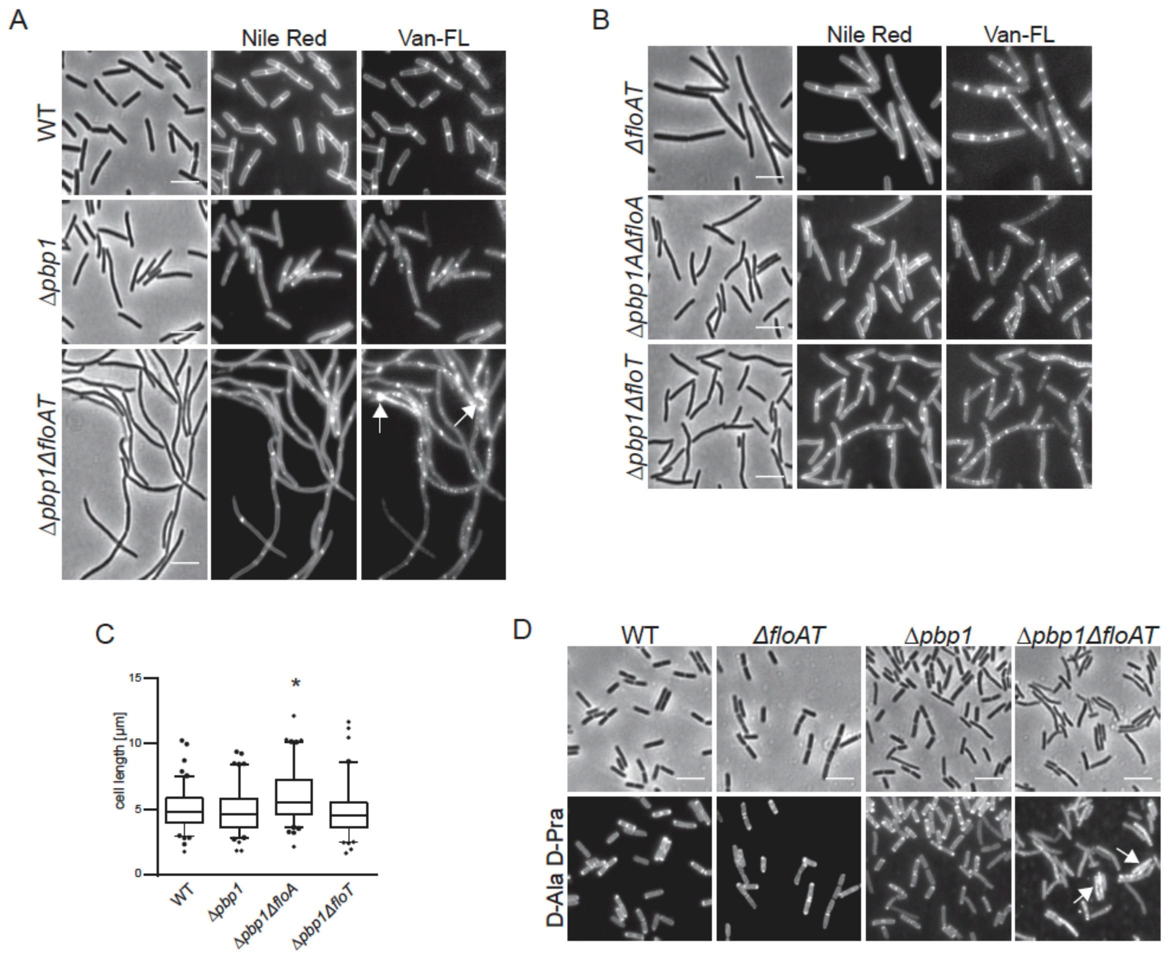


**Figure 2 – Figure supplement 1. Deletion of both flotillins and PBP1 induces filamentation and delocalization of peptidoglycan synthesis.**

**A**. Exponentially growing wt, Δ*pbp1* and Δ*pbp1*Δ*floAT* strains were labelled with Nile Red and Vancomycin-FL (Van-FL). Arrows indicate accumulation of the dye. **B**. Exponentially growing Δ*floAT*, Δ*pbp1*Δ*floA* and Δ*pbp1*Δ*floT* strains were labelled with membrane stain Nile Red and cell wall dye Vancomycin-FL (Van-FL). **C**. Distribution of lengths of cells with a combination of deletions of *pbp1* with either *floA or floT,* imaged in A, B. Statistical analysis and generation of box plots was performed with Prism. Significant differences are based on the two-tailed Mann-Whitney test (n = 100, * $p<0.05$). **D**. Exponentially growing wt, Δ*floAT*, Δ*pbp1* and Δ*pbp1*Δ*floAT* strains were labelled with fluorescent azide bound to D-Ala-D-Pra dipeptide. Arrows indicate accumulation of the dye. There are less filaments observed in this procedure as some filaments break during the fixation procedure that precedes the click reaction. Scale bar (same for all) 5 µm.

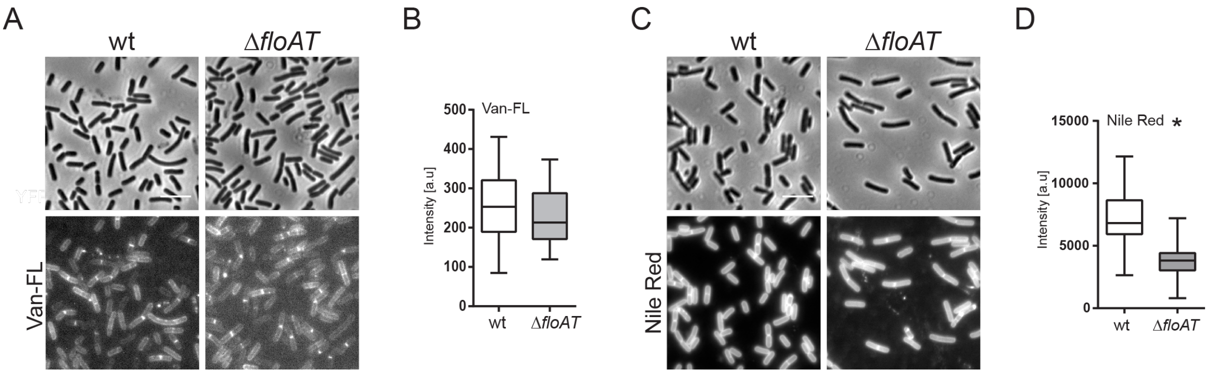


**Figure 2 – Figure supplement 2. Septum labelling of wild type and flotillin mutant cells grown on minimal medium.**

**A, C.** Morphology of the exponentially growing wt and Δ*floAT* strains in the minimal medium labelled with fluorescent Vancomycin (Van-FL, **A**) and Nile Red (**C**). Scale bar 5 µm.

**B, D.** Peak intensity of Van-FL (**B**), and Nile Red (**D**) labelled division sites of the cells shown in (**A** and **C**). Cells from each strain (n=50) were analysed using the ObjectJ macro tool PeakFinder followed by statistical analysis with Prism, where the significant difference is based on an unpaired Welch t-test (* $p < 0.05$).

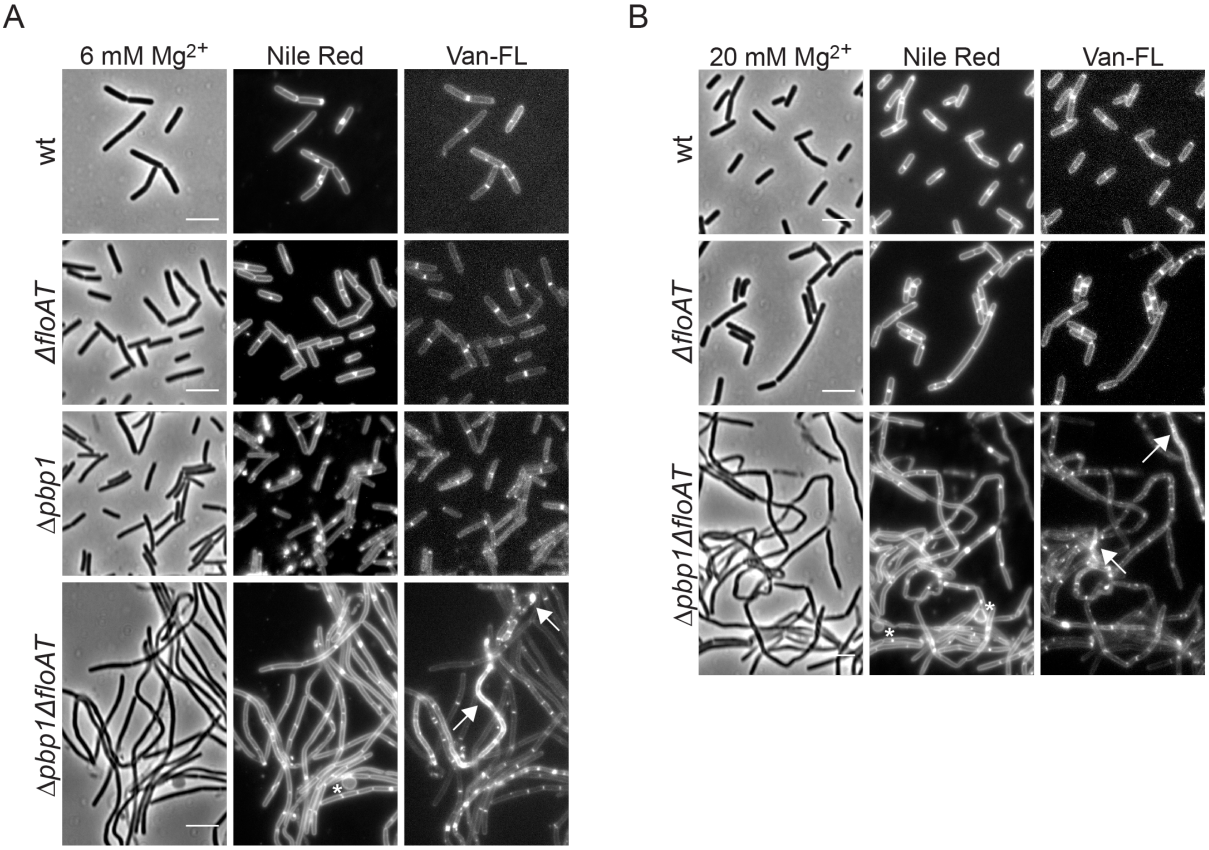


**Figure 2 – Figure supplement 3. Filamentation and delocalization of peptidoglycan synthesis in the absence of flotillins and PBP1 is not rescued by the addition of magnesium.**

Cell morphology of wt, Δ*floAT*, Δ*pbp1*, and Δ*pbp1*Δ*floAT* strains grown in LB supplemented with (**A**) 6 mM magnesium ($Mg^{2+}$) or (**B**) 20 mM magnesium, labelled with Nile Red and Vancomycin-FL (Van-FL). Exponentially growing cells were labelled, and imaged directly with phase contrast and fluorescence microscopy. Arrowheads indicate accumulation of the dye. Scale bar 5 μm.

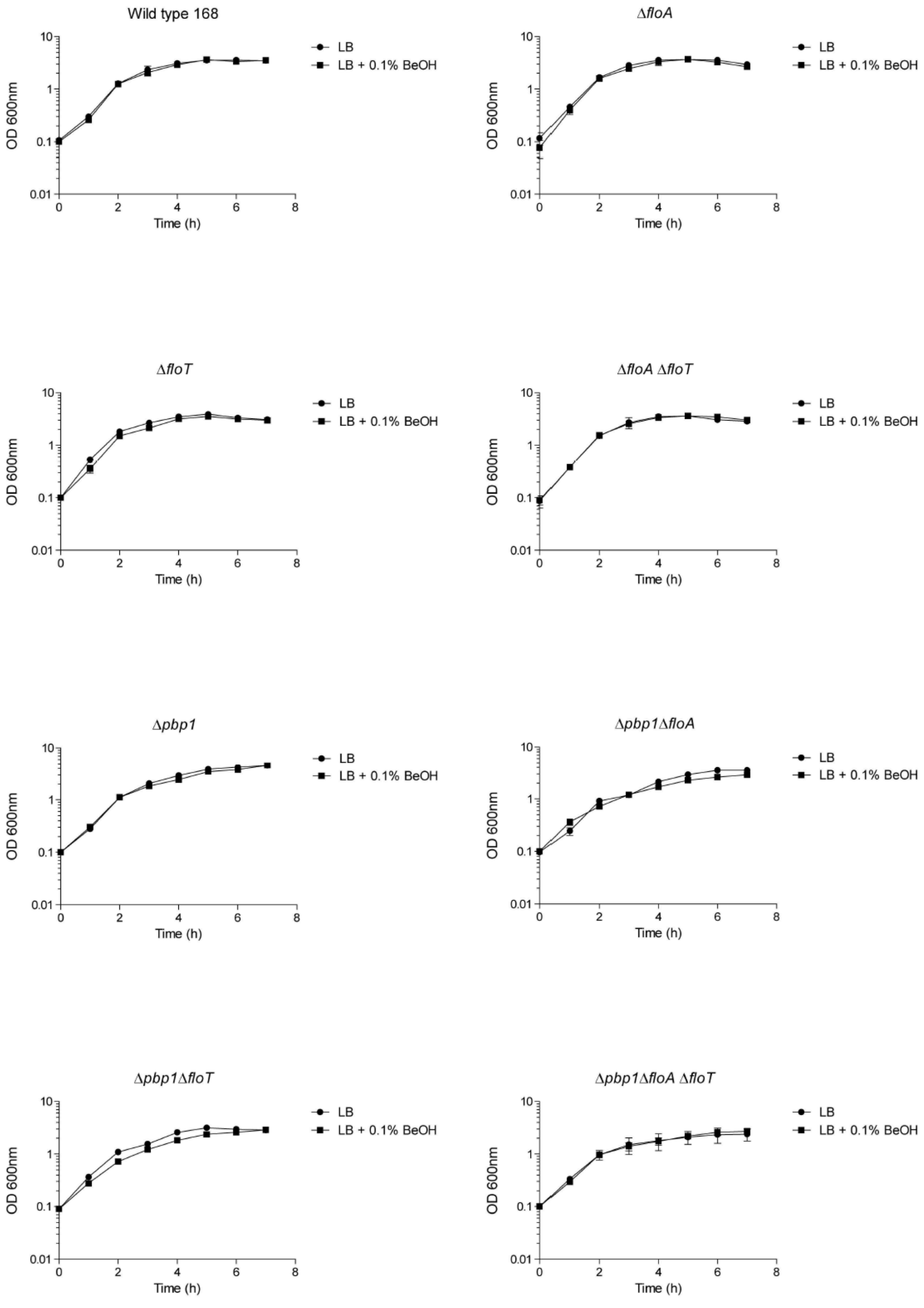


| | μ (h$^{-1}$) | |
|---|---|---|
| strain | LB | LB + 0.1 % BnOH |
| WT | 1.24 ± 0.04 | 1.26 ± 0.03 |
| Δ*floAT* | 1.45 ± 0.15 | 1.42 ± 0.06 |
| Δ*pbp1* | 1.22 ± 0.05 | 1.21 ± 0.03 |
| Δ*pbp1*Δ*floAT* | 1.13 ± 0.11 | 1.13 ± 0.05 |

**Figure 2 – Figure supplement 4. Growth curves and growth rates show similar growth for wt, Δ*floAT*, Δ*pbp1*, and Δ*pbp1*Δ*floAT* (as well as Δ*floA, ΔfloT, Δpbp1ΔfloA and Δpbp1ΔfloT)* strains grown on LB or on LB supplemented with BnOH (0.1 % (w/v)).** Each datapoint represents the average from biological triplicates and error bars indicate standard deviation.

A

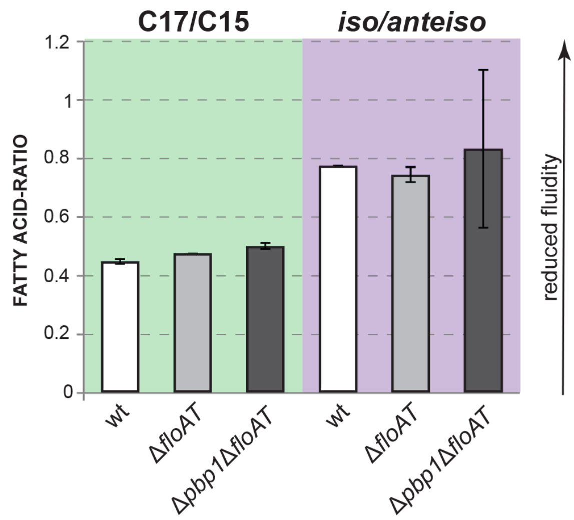


B

| FATTY ACID | wt | Δ*floAT* | Δ*pbp1*Δ*floAT* |
|---|---|---|---|
| $C_{14:0}$ *iso* | 1.6 ±0.07 | 1.3 ±0.07 | 1.8 ±0.48 |
| $C_{15:0}$ *iso* | 20.2 ±0.3 | 19.9 ±0.05 | 19.0 ±4.09 |
| $C_{15:0}$ *anteiso* | 38.1 ±0.76 | 38.3 ±1.28 | 36.6 ±4.32 |
| $C_{16:0}$ *iso* | 5.3 ±0.02 | 4.4 ±0.34 | 5.9 ±1.38 |
| $C_{16:0}$ | 5.4 ±0.77 | 5.2 ±0.34 | 6.2 ±1.25 |
| $C_{17:0}$ *iso* | 12.8 ±0.63 | 13.5 ±0.28 | 14.2 ±1.87 |
| $C_{17:0}$ *anteiso* | 13.4 ±0.32 | 14.2 ±0.3 | 13.7 ±2.53 |

| | | | |
|---|---|---|---|
| C17/C15 | **0.45** ±0.00 | **0.48** ±0.00 | **0.50** ±0.00 |
| *iso*/*anteiso* | **0.77** ±0.00 | **0.74** ±0.02 | **0.83** ±0.27 |

C

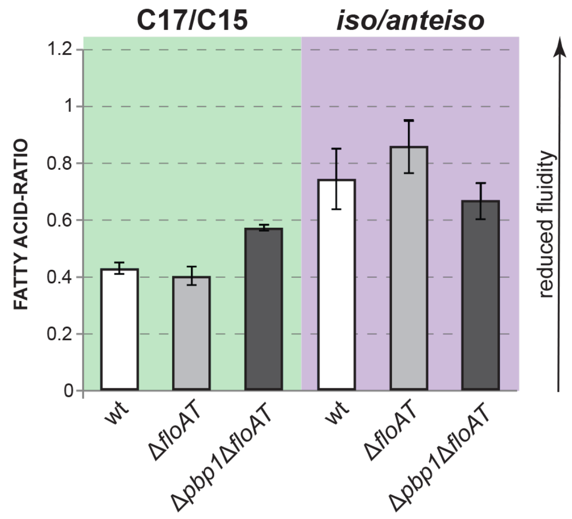


D

| FATTY ACID | wt | Δ*floAT* | Δ*pbp1*Δ*floAT* |
|---|---|---|---|
| $C_{14:0}$ *iso* | 2.6 ±0.16 | 3.1 ±0.8 | 3.3 ±0.14 |
| $C_{15:0}$ *iso* | 19.4 ±3.05 | 21.6 ±1.44 | 11.3 ±1.2 |
| $C_{15:0}$ *anteiso* | 40 ±1.84 | 38.2 ±1.12 | 38.9 ±1.38 |
| $C_{16:0}$ *iso* | 9.2 ±0.82 | 9.7 ±0.96 | 13.6 ±0.19 |
| $C_{16:0}$ | 2.6 ±0.01 | 2.4 ±0.02 | 2.5 ±0.05 |
| $C_{17:0}$ *iso* | 9.9 ±1.12 | 10.2 ±0.53 | 10.1 ±0.7 |
| $C_{17:0}$ *anteiso* | 15.6 ±1.79 | 13.9 ±1.29 | 18.7 ±1.31 |

| | | | |
|---|---|---|---|
| C17/C15 | **0.43** ±0.02 | **0.4** ±0.03 | **0.57** ±0.01 |
| *iso*/*anteiso* | **0.74** ±0.1 | **0.86** ±0.09 | **0.66** ±0.06 |

**Figure 3 – figure supplement 1. Fatty acid composition analysis.**

**A**, **C**. Ratios between chain lengths of the major fatty acids (C17 and C15) and ratios between the *iso* and *anteiso* forms of fatty acids of the exponentially growing wt, Δ*floAT* and Δ*pbp1*Δ*floAT* strains cultivated in LB (A) or minimal medium (C). **B**, **D** - Corresponding total fatty acid profiles and determination of the ratios in the graphs A and C, respectively. Lipid species contributing more that 1% to overall membrane composition are shown. The charts represent the average value of two independent analyses. Fatty acid ratios remain stable upon deletion of flotillins. Triple deletion of both flotillins and PBP1 causes some variability of the total lipid fractions.

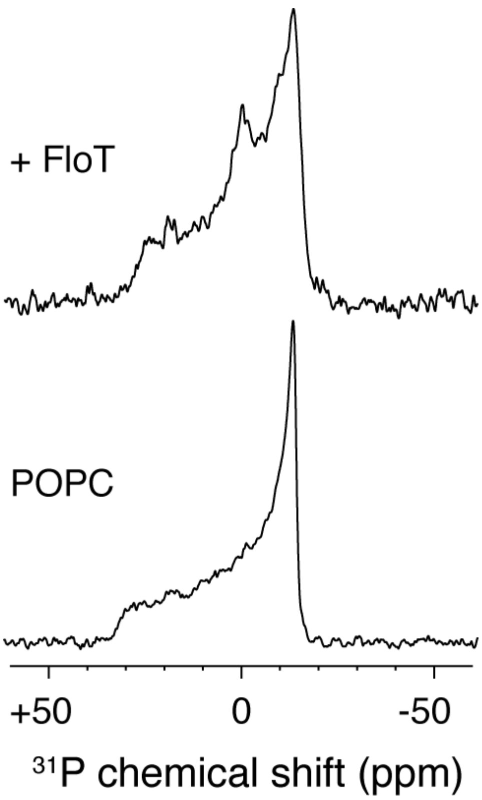


**Figure 5 – Figure supplement 1. $^{31}P$ solid-state NMR experiments of POPC liposomes with or without FloT at a lipid-to-protein molar ratio of 25:1.** All the spectra were acquired at 298 K and a Lorentzian line broadening of 50 Hz was applied before the Fourier transformation.
